# Organic manure managements increases soil microbial community structure and diversity in double-cropping paddy field of Southern China

**DOI:** 10.1101/2020.04.08.031609

**Authors:** Haiming Tang, Xiaoping Xiao, Chao Li, Xiaochen Pan, Kaikai Cheng, Lihong Shi, Ke Wang, Weiyan Li

## Abstract

The soil physicochemical properties were affected by different fertilizer managements, and the soil microbial communities were changed. Fertilizer regimes were closely relative to the soil texture and nutrient status in a double-cropping paddy field of southern China. However, there was limited information about the influence of different long-term fertilizer management practices on the soil microbial communities in a double-cropping rice (*Oryza sativa* L.) fields. Therefore, the 39-year long-term fertilizer regimes on soil bacterial and fungal diversity in a double-cropping paddy field of southern China were studied by using Illumina sequencing and quantitative PCR technology in the present paper. The filed experiment were including chemical fertilizer alone (MF), rice straw residue and chemical fertilizer (RF), 30% organic manure and 70% chemical fertilizer (OM), and without fertilizer input as a control (CK). The results showed that diversity indices of soil microbial communities with application of organic manure and rice straw residue treatments were higher than that without fertilizer input treatment. Application of organic manure and rice straw residue managements increase soil bacterial abundance of the phylum *Proteobacteria, Actinobacteria*, and *Gammaproteobacteria*, and soil fungi abundance of the phylum *Basidiomycota, Zygomycota* and *Tremellales* were also increased. Compared with CK treatment, the value of Richness, Shannon and McIntosh indices, and taxonomic diversity were increased with RF and OM treatments. This finding demonstrated that RF and OM treatments modify soil bacterial and fungal diversity. Therefore, the combined application of organic manure or rice straw residue with chemical fertilizer managements could significantly increase the abundance of profitable functional bacteria and fungi species in double-cropping rice fields of southern China.

## Introduction

Soil microbial plays an important role in soil nutrient cycle, which were affected by different fertilizer regimes and was closely relative with crop growth, soil fertility, and sustainability of soil productivity [1]. Soil microbial activity and community structure were affected by different agricultural managements, such as the crop types, soil tillage, fertilizer regimes, irrigation patterns, and so on. Soil biological properties were mainly affected by different fertilizer management practices, and the soil microbial activities and communities were changed [2]. Maintaining the complexity and diversity of soil microbial is critical to sustain soil fertility, because soil microbes play vital role in mediate the cycles of carbon (C) and nitrogen (N), as well as serve as an important reservoir for plant nutrients [3].

Soil microbial communities exhibit immense diversity in terms of structure (i.e. community composition) as well as function (i.e. physiology of all species combined) that vary from place to place as well as over time. In recent years, the effects of different fertilizer regimes on soil microbial properties were conducted by more and more researchers. Fertilizer treatments affect soil microbial communities through direct influence on soil physical properties and nutrient contents [4]. Organic farming practices (i.e. organic manure and crop residues) generally have positive effects on various soil properties [5], enhancing soil structure by increasing soil organic matter (SOM) content, and thus maintaining soil fertility [6]. Therefore, the influence of fertilizer management practices was provided a way for us to study the soil biological processes. In the previous studies indicated that fertilizer practices was major factor in affecting community structure and diversity of soil microbes, diversity and stability of soil microbial community were enhanced by taken organic input management practices [7]. Some results also indicated that organic manure managements had positive effects on the SOM content, soil microbial community structure and diversity, and the abundance of soil bacteria and fungi [8-9]. Ahn et al. (2016) [10] results indicated that the abundance, activity and diversity of soil microbes were enhanced with return of organic manure and crop residues to soils. In contrast, some study indicated that application of chemical fertilizer management practices increased soil microbial community and activity [11]. However, there is still limited information about the change of soil microbial community structure and diversity under different long-term fertilizer management practices conditions.

Rice (*Oryza sativa* L.) is the major crop in the tropical and subtropical monsoon climate regions of Asia [12]. The early rice and late rice (double-cropping) production system is the mainly crop system in southern of China, and the fertilizer regime (organic manure, crop residues, inorganic fertilizer, and so on) is an important influence factors that maintain the quality and fertility of paddy soils [13]. And the soil properties including the soil pH, soil organic carbon (SOC) content were affected by different long-term fertilizer management practices, which in turn influence the soil bacteria and fungi community structure and function [14-15]. However, there is little information about the influences of 39-year long-term fertilizer regimes on soil microbial activities and communities structure in double-cropping paddy field of southern China. We hypothesized that the function and structure of soil bacteria and fungi communities were changed under taken different fertilization management’s conditions. Therefore, the abundance and community structure of soil bacteria and fungi were investigated by using Illumina sequencing and quantitative quantitative real-time polymerase chain reaction (PCR) technology, respectively.

As a result, the object of this study were: (1) to analyze the effect on soil microbial community following 39-year of continuous application of different fertilization managements, (2) to investigate the relative between soil bacteria, fungi community and soil properties, and (3) to choose an proper fertilizer practice in a double-cropping paddy field of southern China.

## Materials and methods

### Sites and cropping system

The experiment was beginning in 1986. It was located in NingXiang County (28°07′ N, 112°18′ E) of Hunan Province, China. The climatic conditions of the experiment field, the surface soil physical and chemical properties (0–20 cm) beginning of this experiment and crop rotation systems as described by Tang *et al*. (2018) [14].

### Experimental design

The experiment including four treatments: control (without fertilizer input, CK), chemical fertilizer alone (MF), rice straw residue and chemical fertilizer (RF), and 30% organic manure and 70% chemical fertilizer (OM). A randomized block design was adopted in the plots, with three replications. And each plot size was 66.7 m^2^ (10 × 6.67 m). The experiment ensured that the same amount of N, phosphorus pentoxide (P_2_O_5_), potassium oxide (K_2_O) for all fertilizer treatments during the early and late rice growing season, respectively. Details information about the fertilization managements and farming arrangements as described by Tang *et al*. (2018) [14].

### Soil sampling

Soil samples at 0–20 cm depths were collected on October 2019 (after late rice harvest) from each plot. Soil samples were collected by randomly from six cores from each plots, plant roots were removed from the samples and then passing through a 2-mm mesh sieve. The fresh soil samples were placed immediately on ice and transported to the laboratory, and aliquots of the soil samples were then stored at room temperature until soil properties analysis (for pH, SOC, and total nitrogen), at −20°C until molecular analysis.

### Soil laboratory analysis

#### Physical and chemical characteristics

The physical and chemical analyses of the soil were performed in the laboratory. Measurements of pH, SOC, total nitrogen, and soil porosity were performed according to Zhao et al. (2014) [16]. Soil microbial biomass carbon (SMBC) and soil microbial biomass nitrogen (SMBN) contents were measured by the fumigation-extraction method as described in Wu et al. (1990) [17].

#### DNA extraction, PCR amplification, and Illumina sequencing

The concentration and quality of the soil DNA were detected following the manufacturer’s instructions. Primers pairs F515 and R806 targeting the V3-4 region of the soil bacteria 16S rRNA gene were used for PCR [18]. And the protocol for PCR amplification of the 16S rRNA gene were performed using the method described by Wang et al. (2016) [19]. And the target DNA fragment (ITS1 region) of soil fungal was amplified by PCR reaction system with ITS1 and ITS2 as primers [20]. Each PCR product was subjected to pyrosequencing using the Illumina MiSeq platforms at Xidai Biotechnology Co., Ltd., Changsha, China. The original DNA fragments were merging the pairs of reads using FLASH software [19]. Therefore, further sequence analyses were conducted using USEARCH v5.2.32 (Edgar, 2010) [21].

#### Sequence analysis and population identification

The merge and OTU partition of the obtained sequence, and select the representative sequence of each OUT by using QIIME software. Therefore, the default parameters were used to obtain the corresponding taxonomic information of each OTU by comparing the representative sequence of each OTU with the template sequence of the corresponding database. The gene database of soil bacteria was Greenenes v13.8, and the identification of soil fungi was compared with UNITE database (http://unite.ut.ee).

#### Statistical analysis

The Richness diversity, Shannon diversity and McIntosh diversity indices were calculated using Mothur software. Meanwhile, the relationship between abundance of dominant phyla and soil physicochemical properties were measured using redundancy analysis.

The results of every measured item were presented in mean values and standard error. The data of each treatment means were compared by using one-way analysis of variance (Anova) following standard procedures at the 5% probability level. All statistical analyses of correlations between soil properties and abundant phyla were calculated by using the SAS 9.3 software package [22].

## Results

### Soil properties

The effects of different long-term fertilizer management practices on soil pH, soil organic carbon (SOC) content, porosity, total N, soil microbial biomass carbon (SMBC), and soil microbial biomass nitrogen (SMBN) contents in a double-cropping rice field were shown in Table 1. At maturity stage of late rice, the lowest soil pH, SOC, porosity, total N, SMBC, and SMBN in paddy field were observed with MF and CK treatments, while the highest soil pH, SOC, porosity, total N, SMBC, and SMBN contents in paddy field were observed with OM treatment. Meanwhile, the results showed that the SOC, porosity, total N, SMBC, and SMBN contents with OM treatment were significantly higher (*p*<0.05) than that of MF and CK treatments in a double-cropping rice paddy field.

**Table 1.**
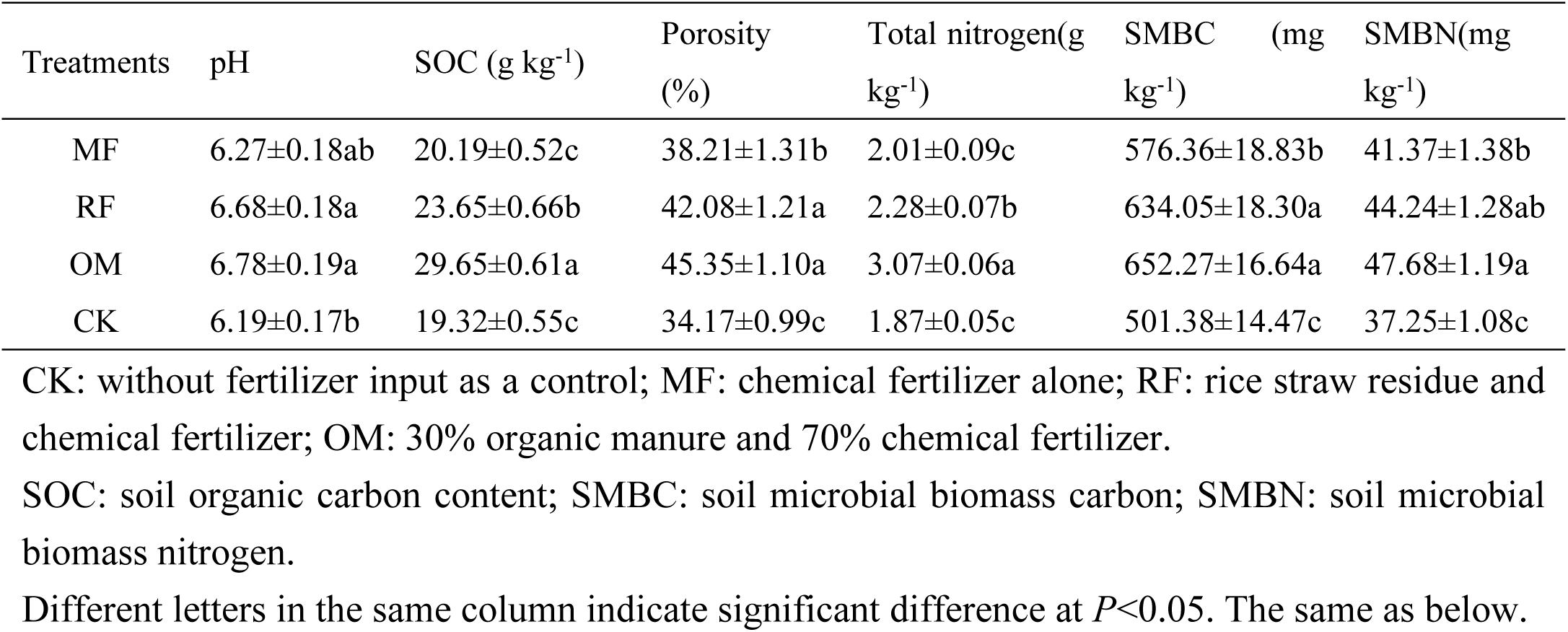
Basic soil properties with different long-term fertilizer treatments in paddy field at maturity stage of late rice

| Treatments | pH | SOC (g kg<sup>-1</sup>) | Porosity (%) | Total nitrogen(g kg<sup>-1</sup>) | SMBC (mg kg<sup>-1</sup>) | SMBN(mg kg<sup>-1</sup>) |
| --- | --- | --- | --- | --- | --- | --- |
| MF | 6.27±0.18ab | 20.19±0.52c | 38.21±1.31b | 2.01±0.09c | 576.36±18.83b | 41.37±1.38b |
| RF | 6.68±0.18a | 23.65±0.66b | 42.08±1.21a | 2.28±0.07b | 634.05±18.30a | 44.24±1.28ab |
| OM | 6.78±0.19a | 29.65±0.61a | 45.35±1.10a | 3.07±0.06a | 652.27±16.64a | 47.68±1.19a |
| CK | 6.19±0.17b | 19.32±0.55c | 34.17±0.99c | 1.87±0.05c | 501.38±14.47c | 37.25±1.08c |

CK: without fertilizer input as a control; MF: chemical fertilizer alone; RF: rice straw residue and chemical fertilizer; OM: 30% organic manure and 70% chemical fertilizer.
SOC: soil organic carbon content; SMBC: soil microbial biomass carbon; SMBN: soil microbial biomass nitrogen.
Different letters in the same column indicate significant difference at $P<0.05$ . The same as below.

### Soil bacterial taxonomic distribution

The phylum level analysis results indicated that phyla *Actinobacteria, Acidobacteriales, Alphaproteobacteria, Chloroflexi, Fimicutes, Proteobacteria* and *Planctomycetes* were the most abundant in the soil samples. And the phyla *Proteobacteria, Actinobacteria* with RF and OM treatments were significantly higher than that of with FM and CK treatments (Fig. 1). The phyla *Gemmatimonadetes* and *Acidobacteriales* with RF and OM treatments were significantly lower than that of with CK treatment. The results also indicated that relative abundances of *Gammaproteobacteria* were richer higher with RF and OM treatments soils, and the relative abundances of *Fimicutes, Alphaproteobacteria, Planctomycetes* and *Chlorobi* were higher with CK treatment soils. Significant differences in soil bacterial composition were observed between the RF, OM and FM, CK treatments soils (Fig. 1).

**Fig. 1.**
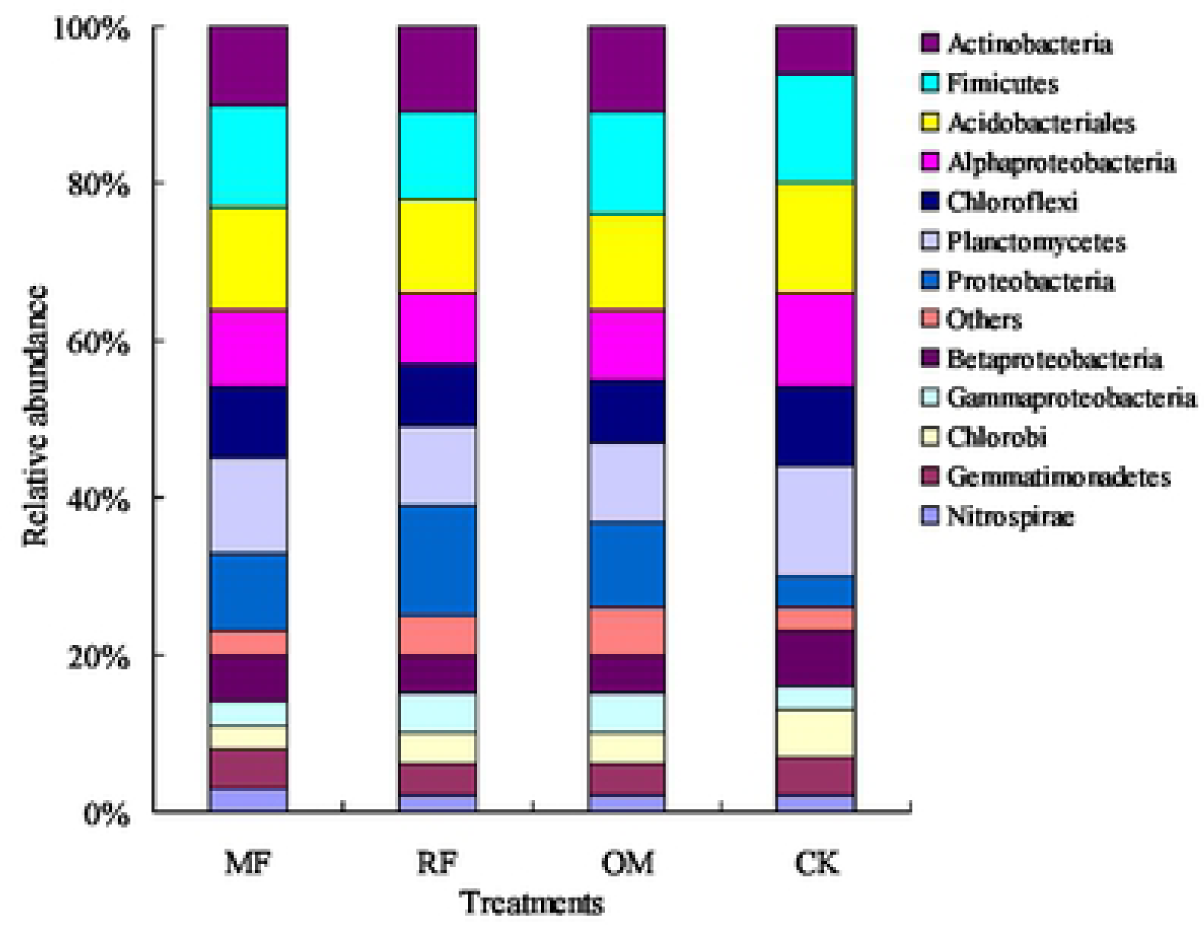
Relative abundance of the dominant bacterial phyla in all soil samples combined and in each long-term fertilizer treatments CK: without fertilizer input as a control; MF: chemical fertilizer alone; RF: rice straw residue and chemical fertilizer; OM: 30% organic manure and 70% chemical fertilizer. Relative abundances were based on the proportional frequencies of those DNA sequences that could be classified. The same as below.

### Soil fungal taxonomic distribution

The results showed that phyla *Ascomycota, Sordariales, Basidiomycota* and *Zygomycota* were the most abundant in the soil samples. And the order of phyla with different fertilizer treatments was showed *Ascomycota*>*Basidiomycota*>*Sordariales*>*Zygomycota*. And the phyla *Ascomycota* with CK treatment were significantly higher than that of with MF, RF and OM treatments (Fig. 2). The phyla *Basidiomycota* and *Zygomycota* with RF and OM treatments were significantly higher than that of with MF and CK treatments. The results indicated that relative abundances of *Basidiomycota, Zygomycota* and *Tremellales* were richer higher with RF and OM treatments soils, and the relative abundances of *Ascomycota, Hypocreales*, and *Pleosporales* were higher with CK treatment soils.

**Fig. 2.**
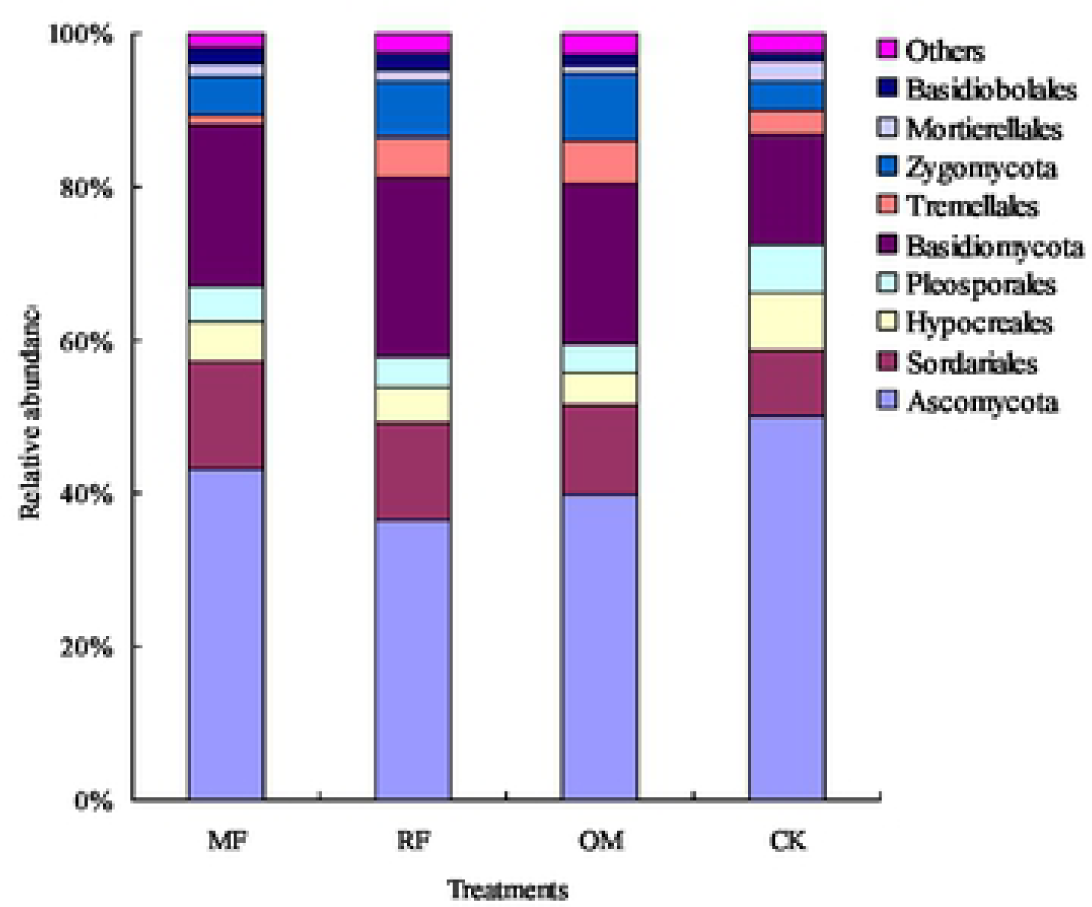
Relative abundance of the dominant fungal phyla in all soil samples combined and in each long-term fertilizer treatments.

### Soil bacterial and fungi alpha diversity

Richness, Shannon and McIntosh indices were used to reflect the richness and evenness of soil microbial community species, respectively. At soil bacterial alpha diversity, Richness indices were increased with OM treatment compared with CK treatment, and the order of was OM>RF>MF>CK. Compared with CK treatment, the Shannon and McIntosh indices were increased with MF, RF and OM treatments (Table 2). At soil fungi alpha diversity, Richness indices were increased with OM treatment compared with CK treatment. Compared with CK treatment, the Shannon and McIntosh indices were increased with RF and OM treatments (Table 2). The results indicated that Richness indices, Shannon indices and McIntosh indices with application of rice straw residue and organic manure managements were higher than that of with chemical fertilizer alone and without fertilizer input managements.

**Table 2.** Soil bacterial and fungal diversity parameters with different long-term fertilizer treatments

| Microorganism | Diversity parameters | Treatments |  |  |  |
| --- | --- | --- | --- | --- | --- |
|  |  | MF | RF | OM | CK |
| Soil bacterial | Richness indices | 14.61±0.41b | 15.02±0.42ab | 15.35±0.42a | 14.28±0.41b |
|  | Shannon indices | 4.84±0.16b | 5.26±0.15ab | 5.63±0.14a | 4.17±0.12c |
|  | McIntosh indices | 5.85±0.18b | 6.14±0.18ab | 6.48±0.17a | 4.73±0.14c |
| Soil fungi | Richness indices | 11.03±0.34ab | 11.35±0.33ab | 11.96±0.32a | 10.74±0.31b |
|  | Shannon indices | 3.72±0.12b | 4.06±0.12ab | 4.17±0.11a | 3.21±0.10c |
|  | McIntosh indices | 4.51±0.14ab | 4.63±0.13a | 4.85±0.13a | 4.19±0.12b |

Different letters in the same line indicate significant difference at $P<0.05$ . The same as below.

### RDA of soil bacterial and fungal abundant phyla and soil characteristics

The results revealed there was a significant correlation between soil bacterial abundant phyla and soil chemical properties (Fig. 3 a). The soil properties can explain the variation (85.78% and 15.46%) in soil bacterial abundant phyla between the MF, RF, OM and CK treatments. Under different long-term fertilizer treatments, MF and CK treatments were separated from application of rice straw residue and organic manure (RF and OM) treatments, indicating that fertilizer treatments significantly changed the characteristics of soil bacterial abundant phyla. The soil bacterial abundant phyla was significantly correlated with the soil physical and chemical properties including the porosity (*r*=0.894), SOC (*r*=0.916), total nitrogen (*r*=0.904), SMBC (*r*=0.902), and SMBN (*r*=0.906) contents. But there was negative correlated between soil bacterial abundant phyla and pH (*r*= −0.805).

**Fig. 3.**
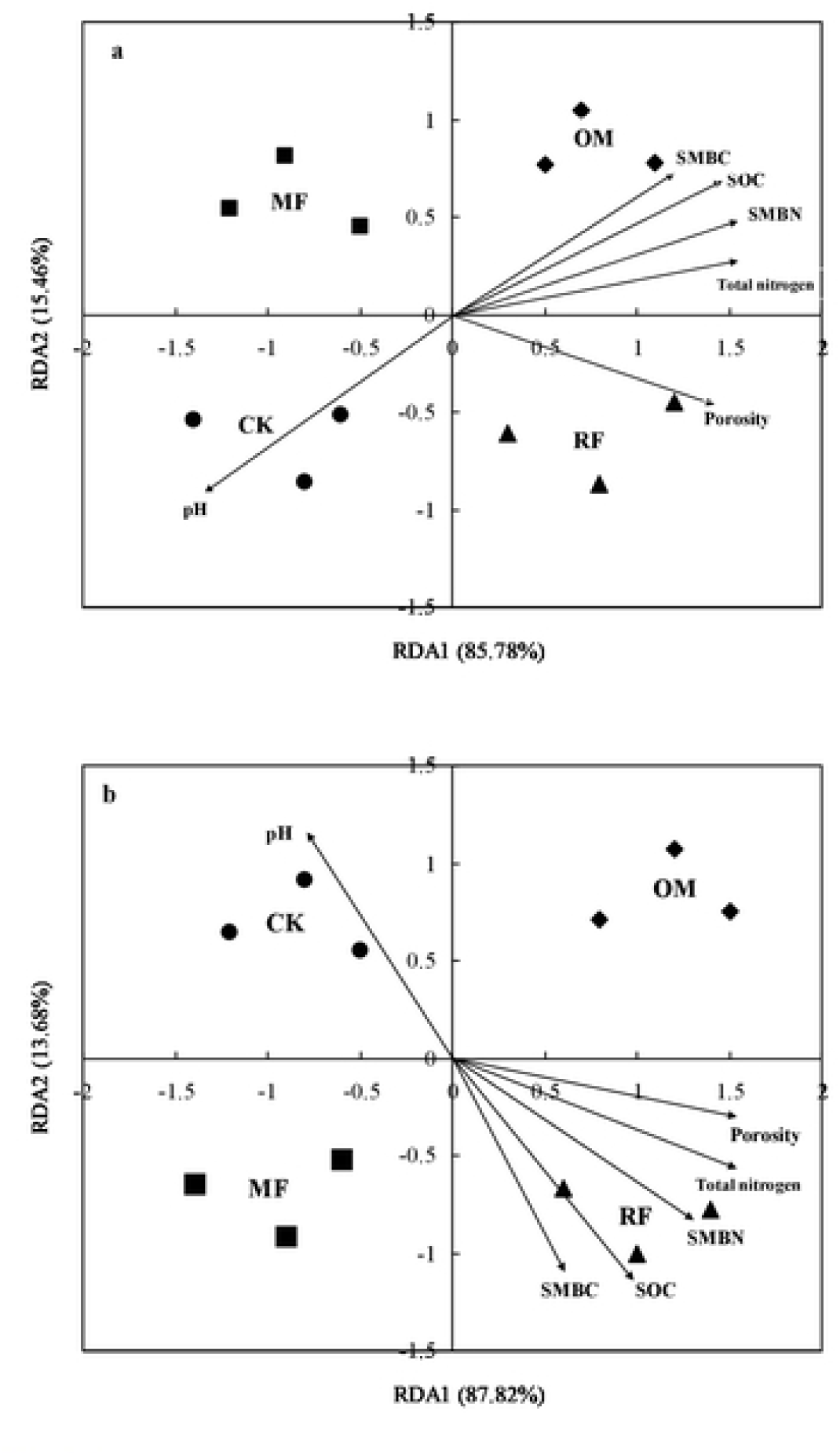
Redundancy analysis of soil bacterial (a) and fungal (b) abundant phyla and soil characteristics (arrows) of different long-term fertilizer treatments CK: without fertilizer input as a control (circle); MF: chemical fertilizer alone (square); RF: rice straw residue and chemical fertilizer (triangle); OM: 30% organic manure and 70% chemical fertilizer (prismatic).

The results indicated that the soil properties can explain the variation (87.82% and 13.68%) in soil fungal abundant phyla between the MF, RF, OM and CK treatments (Fig. 3 b). Under different long-term fertilizer treatments, MF and CK treatments were separated from application of rice straw residue and organic manure treatments (RF and OM treatments). The soil fungal abundant phyla was significantly correlated with the soil physical and chemical properties including the porosity (*r*=0.794), SOC (*r*=0.886), total nitrogen (*r*=0.845), SMBC (*r*=0.724), and SMBN (*r*=0.806) contents. But there was negative correlated between soil fungal abundant phyla and pH (*r*= −0.735).

## Disscussion

In the present study, the results showed that soil microbial community were mainly affected and changed by different long-term fertilizer management practices, including soil structure, soil pH, SOC, total nitrogen, SMBC, and SMBN contents (Table 1), the soils total nitrogen, SOC, SMBC, and SMBN contents were increased with application of organic manure and crop residues incorporation treatments compared with the chemical fertilizer alone and without fertilizer input treatment soils, consistent with the results of a previous study [1, 4]. Application of organic manure and crop residues practices significantly increased the SOC content, the reason maybe that soil C redistribution and microbial habitat were altered under manure and crop residues input managements conditions [23]. Soil aggregation mediated were changed by porosity, and soil C sequestration were increased under application of organic manure and crop residues conditions [24]. Based on those findings, the results were indicated that distribution of the soil bacterial and fungal communities in a double-cropping rice system were changed due to the changes in soil properties.

Compared to the chemical fertilizer alone and without fertilizer input (MF, CK) treatments, the organic matter incorporation (RF and OM) treatments strongly increased the Richness, Shannon and McIntosh indices in the soil samples. At the indices of soil microbial communities, the results indicated that the indices of soil microbial communities were increased with application of organic manure and crop residues management practices (Fig. 1, Fig. 2), which may be the disturb of microbiota were moderated, the soil texture and moisture content were increased, therefore, the competitive niche and selection mechanisms among different populations were excluded. As a result, the Richness, Shannon and McIntosh indices were increased in agree with the previous study [1, 4, 6]. This result maybe explained from the soil texture and moisture conditions using organic matter (organic manure and crop residues) management practices. The soil pore connectivity were mainly affected by soil texture and moisture, which was the important influence factor of changes in soil bacterial and fungal diversity, and was closely related to the Simpson and Shannon indices [7]. There were more nutrient and C source with organic manure and crop residues management practices than that with chemical fertilizer and without fertilizer input treatments, which was provide an appropriate soil environment and nutrient for soil bacteria and fungal multiplying. Therefore, this study was supports the idea that soil bacteria and fungal alpha diversity and abundance were changed with application of organic manure and crop residues management practices, which the soil texture, soil moisture content, and soil properties were obvious altered.

Some studies showed that the structure and diversity of soil bacterial community were changed under different fertilizer conditions [4, 10, 11]. In the present study, the relative abundance of *Proteobacteria* and *Actinobacteria* with RF and OM treatments were higher than that with MF and CK treatments (Fig. 1), which were provide an appropriate soil environment and nutrient for *Proteobacteria* and *Actinobacteria* multiplying under application of organic manure and crop residues conditions. The relative abundance of *Fimicutes* with CK treatment were higher than that with MF, RF and OM treatments, which were mainly benefit for produce endospores [16]. Jenkins et al. (2010) [25] reported that *Proteobacteria* was a fast-growing copiotrophs under higher SOC environments, competed with soil nutrients with *Firmicutes* was the main reason for the reduced abundances of *Alphaproteobacteria* and *Betaproteobacteria* with RF and OM treatments compared to MF and CK treatments soils as observed in the present study. Meanwhile, the results indicated that the abundances of *Alphaproteobacteria* with MF and CK treatments soils were higher than that of the organic matter incorporation (RF and OM) treatments soils, which were considered as heterotrophic, nitrogen-fixing organisms [26]. These findings were significantly correlated with soil texture and soil nutrient conditions change (Table 1), which was main reason for the higher relative abundance of *Alphaproteobacteria* with MF and CK treatments soils than that of with organic matter incorporation treatments soils. Some study indicated that abundance of *Acidobacteriales* and *Chloroflexi* were regarded as oligotrophs [27], and its soil textures and soil nutrient contents were the main reason for the lower abundance of *Acidobacteriales* and *Chloroflexi* with RF and OM treatments soils than that with MF and CK treatments soils. The results indicated that the relative abundances of *Planctomycetes* and *Chlorobi* were higher with CK treatment soils, which was main reason for the growth of *Planctomycetes* and *Chlorobi* were inhibited by the higher SOC contents under application of organic manure and crop residues conditions, were also consistent with previous observations that these phylum were suppressed by organic matter incorporation (RF and OM) treatments [28, 29]. Therefore, fraction of soil texture were changed, soil aeration and nutrient contents were improved with organic matter incorporation (RF and OM) treatments, thus the relative abundance of profitable functional bacteria species were increased.

There were significantly differences in response patterns of soil fungal community structure and diversity to different long-term fertilizer regimes. In this study, the results showed that phyla *Ascomycota, Sordariales, Basidiomycota* and *Zygomycota* were the most abundant in the soil samples, consistent with the results of previous study [30]. And the phyla *Ascomycota* with organic manure and crop residues treatments were significantly lower than that of with CK treatment (Fig. 2), which were considered as the growth of *Ascomycota* was not good with application of organic manure and crop residues management practices. The main reason maybe that it was not benefit for the growth and reproduction of *Ascomycota* under application of organic manure and crop residues conditions. on the other hand, the application of organic manure and crop residues contains a lot of nitrogen, phosphorus and potassium nutrients, and the excessive application of organic manure and crop residues in the soils will promote the pathogenic characteristics of fungi [31], leading to the inhibition of *Ascomycota* growth. Compared with MF and CK treatments, the phyla *Basidiomycota* and *Zygomycota* were increased with RF and OM treatments, which was provide nutrient and appropriate soil environment for *Basidiomycota* and *Zygomycota* multiplying. The reason may be that the application of organic manure and crop residues treatments can increase the SOC content, improve the ventilation condition of soil, and create favorable soil conditions for *Basidiomycota* and *Zygomycota* growth. However, the application of chemical fertilizer alone treatment were cause the deterioration of soil physical and chemical properties and soil hardening, which was not benefit for the growth and reproduction of *Basidiomycota* and *Zygomycota*.

In the present study, the redundancy analysis results indicated that soil bacterial community structure was closely related to soil porosity, SOC, total nitrogen, SMBC, and SMBN contents, and soil fungal community structure was closely related to soil porosity, total nitrogen, SMBN and SOC contents. Meanwhile, there were also obvious effects of soil pH on the soil bacteria and fungi community structure (Fig. 3). Many studies have shown that soil microbial growth was affected by soil texture and soil properties conditions [3]. SMBC content was also an important parameter reflecting the turnover of soil organic matter [32], which was influence on soil bacterial and fungal community structure. Meanwhile, some studies also indicated that soil bacterial and fungal community structure were affected by soil organic matter and total nitrogen contents [8-9]. In this study, the soil fertility were improved with application of organic manure and crop residues management practices, and the soil microorganisms growth and reproduction, carbon/nitrogen content of soil microbial biomass were also increased [33], and soil microbial community structure were changed. Some studies results showed that soil pH has a profound impact on the soil bacterial and fungal community structure [34], and our research results were also indicated that there were obvious effects of soil pH on the soil bacterial and fungal community structure, the reason maybe that the change of soil pH was sufficient impact on the soil microbial community (Table 1). Therefore, the soil microbial function was an important indicator of soil quality, which were influenced by different long-term fertilizer management practices. And the mechanism of soil microbial function responds to different fertilizer management practices needs further study.

## Conclusions

In this study, the results indicated that soil physical and chemical properties were significantly changed under 39-year long-term fertilizer treatments conditions, and the soil bacterial and fungal diversities and community compositions were also modified in a double-cropping paddy field of southern China. And the higher value of Richness, Shannon and McIntosh indices were found in organic manure and rice straw residue soils, and more diversification soil bacterial community were observed in combined application of organic manure or rice straw residue with chemical fertilizer soils, which had higher rich functional microorganisms (including *Proteobacteria, Actinobacteria*, and *Gammaproteobacteria*), and more diversification soil fungi community were observed in combined application of organic manure or rice straw residue with chemical fertilizer soils, which had higher rich functional microorganisms (including *Basidiomycota, Zygomycota* and *Tremellales*). Most importantly, the results demonstrated that fraction of soil physical and chemical properties (porosity, SOC, total nitrogen, SMBC, and SMBN contents) were improved under combined application of organic manure or rice straw residue with chemical fertilizer conditions, leading to changes of the soil bacterial and fungi community. Therefore, it was found that combined application of organic manure or rice straw residue with chemical fertilizer managements is feasible managements contribute to increase soil bacterial and fungi community and to build stable soil environment in a double-cropping paddy field of southern China.

## Acknowledgements

This study was supported by National Natural Science Foundation of China (31872851), Innovative Research Groups of the Natural Science Foundation of Hunan Province (2019JJ10003).

